# The origin of the odorant receptor gene family in insects

**DOI:** 10.1101/259424

**Authors:** Philipp Brand, Hugh M. Robertson, Wei Lin, Ratnasri Pothula, William E. Klingeman, Juan Luis Jurat-Fuentes, Brian R. Johnson

## Abstract

The sense of smell enables the detection and discrimination of airborne chemicals via chemosensory receptors that have evolved independently multiple times throughout the tree of life. In insects, the odorant receptor (OR) gene family is the major chemosensory gene family involved in olfaction and its origin has been hypothesized to coincide with the evolution of a terrestrial lifestyle in hexapods. Missbach et al. (2014) challenged this view and suggested that ORs evolved with an ancestral OR co-receptor (Orco) after the origin of terrestriality, hypothesizing that the OR/Orco system is an adaptation to winged flight in insects instead. Building upon this work, we investigated the genomes of basal hexapod and insect lineages including Collembola, Diplura, Archaeognatha, Zygentoma, Odonata, and Ephemeroptera in an effort to identify the origin of the insect OR gene family. While absent from all non-insect hexapod lineages analyzed, ORs are present in all insect genomes. Orco is absent only in the most ancient insect lineage Archaeognatha. A fully functional OR/Orco system was present in our newly generated genome data of the Zygentoma *Thermobia domestica*. We suggest that ORs did evolve as adaptation to a terrestrial lifestyle outside high-humidity habitats, and not winged flight, representing a key evolutionary novelty in the ancestor of all insects. The OR family is therefore the first known molecular synapomorphy for the Class Insecta.

## Introduction

From bacteria to mammals, living organisms of all levels of complexity have evolved chemosensory receptors to detect and discriminate chemicals in the environment (Wuichet and Zhulin, 2010; Hansson and Stensmyr, 2011). The largest metazoan gene families for example encode tens to hundreds of odorant receptors (ORs) that interact with volatile chemicals at the sensory periphery underlying the sense of smell (Sánchez-Gracia et al., 2009; Niimura et al., 2014). OR gene families have evolved multiple times throughout the metazoans, including independent origins in vertebrates, nematodes, and insects (Hansson and Stensmyr, 2011). In insects, the OR gene family evolved from within the ancestral gustatory receptor (GR) gene family (Scott et al., 2001; Robertson et al., 2003) that extends back to ancient metazoan lineages (Robertson, 2015; Saina et al., 2015; Eyun et al., 2017). ORs are absent from non-insect arthropod genomes (Peñalva-Arana et al., 2009; Almeida et al., 2015; Gulia-Nuss et al., 2016; Ngoc et al., 2016; Eyun et al., 2017), and have been hypothesized to have evolved concomitant with the evolution of terrestriality in hexapods (Robertson et al., 2003).

The lack of molecular resources for early branching hexapod and insect lineages has prevented the precise dating of the origin of insect ORs. Only recently, whole-genome sequencing efforts suggested that ORs are absent in non-insect hexapods such as Collembola (Wu et al., 2017) but present in early branching pterygote insects such as damselflies (Odonata; Ioannidis et al., 2017). Efforts to understand more precisely the origin of the OR gene family within hexapods were greatly advanced by the findings of Missbach et al. (2014) who sequenced transcriptomes of the chemosensory organs of two apterygote insects, the bristletail *Lepismachilis y-signata* (Archaeognatha) and the firebrat *Thermobia domestica* (Zygentoma). They identified three ORs in the firebrat, which they named TdomOrco1–3, with apparent similarity to the neopteran odorant receptor co-receptor (Orco; Vosshall and Hansson, 2011). Orco is a highly conserved single-copy gene present in all other insects studied to date and encodes a protein that is a partner with each of the other “specific” ORs (Benton et al., 2006) which is required for OR-based olfaction in insects (Larsson et al., 2004). In contrast, Missbach et al. (2014) could not find ORs or Orco relatives in their bristletail transcriptome, instead finding only members of the ionotropic receptor (IR) gene family. Given evidence that IRs serve olfactory roles in terrestrial crustaceans and insects (Rytz et al., 2013; Groh-Lunow et al., 2015; Rimal and Lee, 2018), they argued that olfaction in early-branching terrestrial hexapods and apterygote insects is entirely IR-dependent, with Orco evolving as ancestral OR from the GR lineage between the Archaeognatha and Zygentoma. Based on these findings, Missbach et al. (2014) suggested that the Orco/OR system evolved together with flight in pterygote insects and left off with the observation that “the existence of three Orco types remains mysterious”.

Recently, phylogenetic analysis of the OR gene family of the damselfly *Calopteryx splendens* suggested that at least one of the three Orco-like ORs from *T. domestica*, TdomOrco3, might be a specific OR instead of an Orco (Ioannidis et al., 2017). If this is correct, then the entire Orco/OR system evolved before winged insects, which would explain the “mystery” of three apparent Orco types in Zygentoma. In an effort to identify the origin of the insect OR gene family and the Orco/OR system, we investigated the genome sequences of species belonging to multiple ancient terrestrial hexapod and insect orders, including Collembola (springtails), Diplura (two-pronged bristletails), Archaeognatha (jumping bristletails), Zygentoma (silverfish and firebrats), Odonata (damselflies and dragonflies), and Ephemeroptera (mayflies).

## Results and Discussion

### ORs were present in the ancestor of insects

We detected no ORs in two non-insect hexapod lineages, Collembola and Diplura (a genome sequence is not available for the third lineage, the Protura), despite extensive annotation efforts. In contrast, we identified genes with similarity to known insect ORs in all other genomes investigated (Figure 1; details in Supplemental Material, Tables S2 and S3). These included one species of each of the earliest branching insect lineages, the Archeognatha, Zygentoma, Ephemeroptera, and Odonata (Misof et al., 2014). Accordingly, ORs were likely present in the ancestor of all insects but absent from all non-insect hexapod lineages. This suggests that the origin of the OR gene family coincided with the evolution of insects. Thus, our analysis does not support the hypothesis that ORs evolved with the evolution of winged flight in insects (Missbach et al., 2014) but is compatible with the hypothesis that they evolved with terrestriality in insects (Robertson et al. 2003). Terrestrial hexapod lineages without ORs (i.e. Diplura and Collembola) are confined to highly humid environments including moist soil and leaf-litter. This is analogous to terrestrial crustaceans, which evolved OR-independent modified sensory systems allowing the detection of airborne odors on land (Stensmyr et al., 2005; Hansson et al., 2010). While terrestrial crustaceans employ the same olfactory gene-families as their marine ancestors in combination with anatomical adaptations to terrestrial olfaction, their sense of smell is highly dependent on humidity (Krång et al., 2012). Although the molecular and physiological basis of dipluran and collembolan olfaction is unexplored, they are known to have a highly sensitive sense of smell used in pheromone communication and foraging behavior (Verhoef et al., 1977; Purrington et al., 1991; Staaden et al., 2011). It is thus possible that ORs evolved as an adaptation to a terrestrial lifestyle outside ancestrally humid environments in insects.

**Figure 1.**
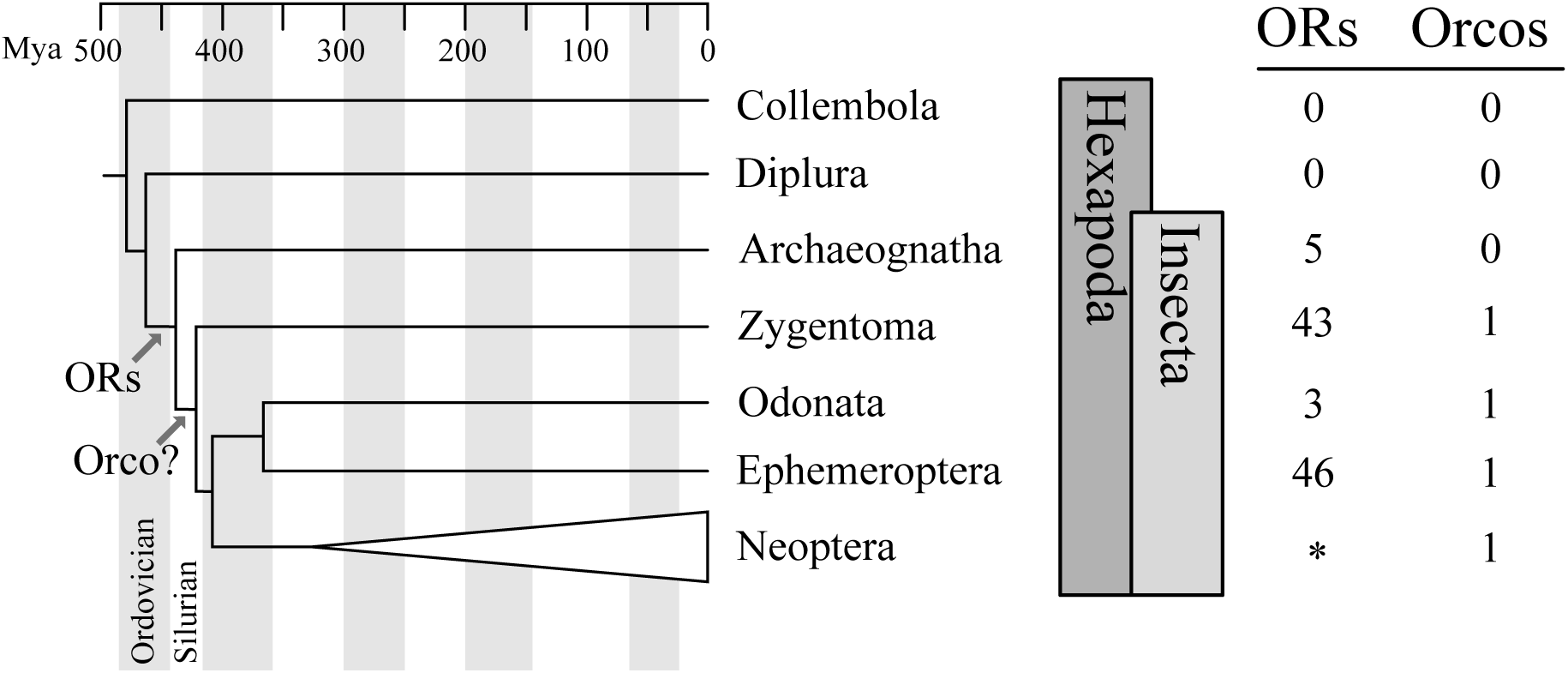
Origin of the insect odorant receptor gene family. The number of ORs and OR co-receptors (Orcos) for all ancient insect and non-insect orders analyzed was mapped on the hexapod phylogeny sensu Misof et al. (2014). ORs are present in all insects but absent from non-insect hexapod genomes, and thus likely represent a molecular synapomorphy for the Clade Insecta. Orco is present in all but Archaeognatha, the earliest branching insect order. This suggests two scenarios including either the loss of Orco in Archaeognatha or an Orco origin following the evolution of ORs (as indicated). The OR gene family likely evolved as an adaptation to a terrestrial lifestyle outside ancestrally humid environments in insects. *: The genomes of all neopteran insects analyzed to date encode ORs, ranging from 10 ORs in head lice (Kirkness et al., 2010) to more than 300 ORs in ants (Smith et al., 2011a; b).

### The *Thermobia domestica* genome harbors a full Orco/OR gene family repertoire

With the exception of the Zygentoma, all lineages analyzed had genome data either published (Faddeeva-Vakhrusheva et al., 2016; 2017; Wu et al., 2017) or available from the i5k Pilot Project from the Baylor College of Medicine at the i5k NAL Workspace (Poelchau et al., 2015). To complete taxon sampling of early branching hexapod and insect lineages, we produced a draft genome assembly for *T. domestica* (Supplemental Material; Figure S1; Table S1). This enabled direct comparison to Missbach et al. (2014) and revealed that the *T. domestica* genome encodes far more than the three Orco-like OR proteins. Our manual annotation revealed 43 ORs encoded by 31 genes including the three previously identified genes (TdomOrco1–3; Missbach et al., 2014). Four genes are modeled as exhibiting alternative splicing leading to the additional protein isoforms (Supplemental Material). We used the antennal transcriptome of Missbach et al. (2014) for support of intron-exon boundaries, however only a few transcriptome reads mapped to the “specific” OR genes (Table S2), indicating that the RNA-seq analysis of Missbach et al. (2014) did not sequence to a sufficient depth to reconstruct these low-expressed transcripts.

Phylogenetic analyses of all ORs we annotated in the bristletail *Machilis hrabei* (5 ORs), the dragonfly *Ladona fulva* (4 ORs), and the mayfly *Ephemera danica* (47 ORs), as well as the previously annotated damselfly *C. splendens* (6 ORs; Ioannidis et al., 2017) revealed that one of the *T. domestica* ORs (TdomOrco2) clustered confidently with the Orco lineage in pterygote insects (Figure 2; Figure S2). We believe this is the sole Orco relative because it shares unique features with the pterygote Orco proteins, such as a TKKQ motif in the expanded intracellular loop 2 (positions 327–330 in DmelOrco), and so we simply rename it TdomOrco. TdomOr1–8 represent a set of Orco-like proteins that share a common gene structure with TdomOrco, with introns in phases 0-2-0-0-0. These last four introns are present in all the other TdomOr genes, as well as those of the bristletail, Odonata, and mayfly, and correspond to the four introns identified by Robertson et al. (2003) as being ancestral to the OR family. The first phase-0 intron of Orco and Or1–8 is the only additional intron shared by most pterygote Orco genes.

**Figure 2.**
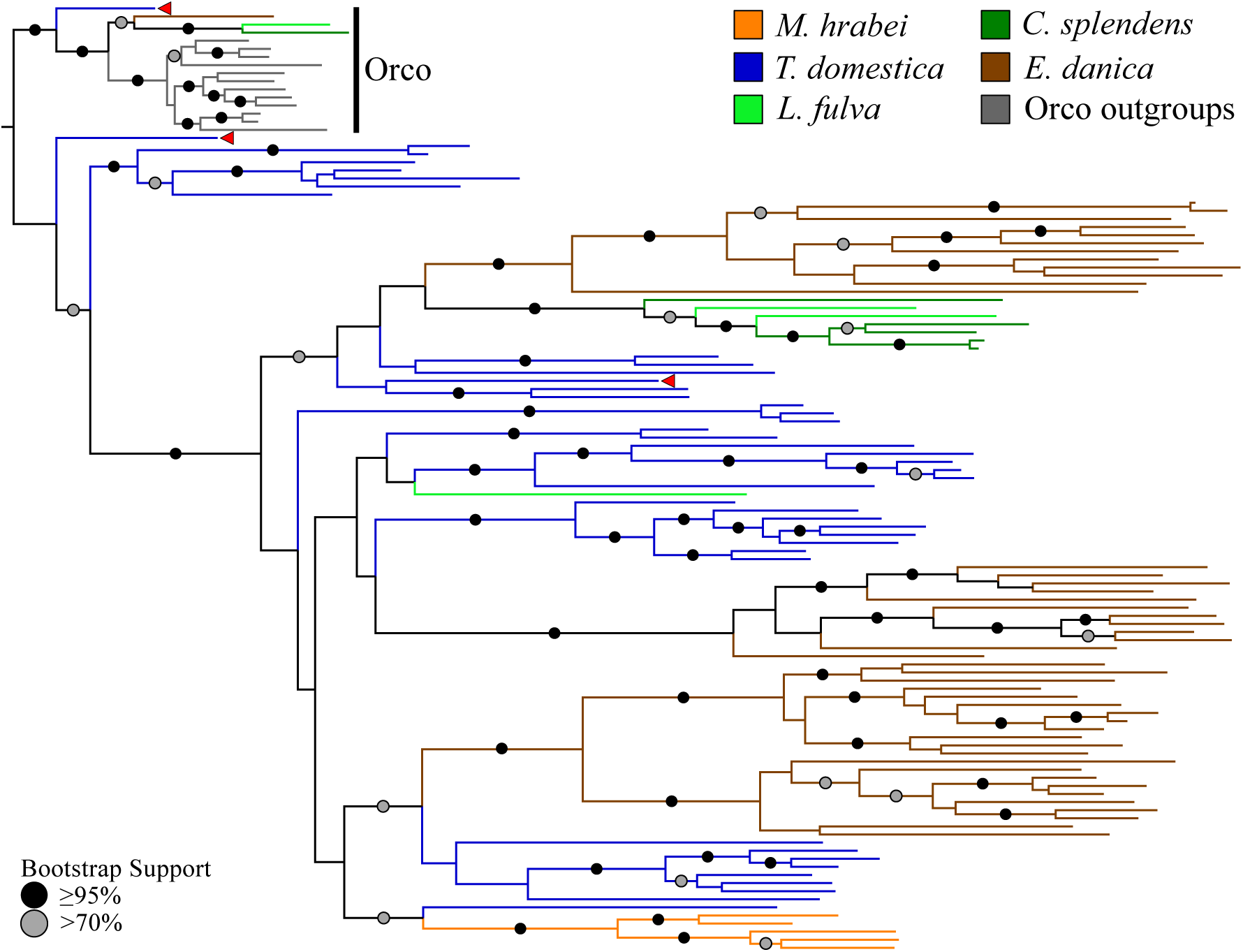
Odorant receptor (OR) gene family phylogeny including representatives of all apterygote and paleopteran insect orders. The Maximum Likelihood tree demonstrates monophyly of the single-copy insect Orco with high bootstrap support. The *M. hrabei* genome lacks Orco but encodes five highly similar ORs clustering in a single highly-supported clade. *T. domestica* has a fully developed functional OR/Orco system. The red arrowhead indicates the location of the three *T. domestica* ORs identified by Missbach et al. (2014).

With the exception of the bristletail *M. hrabei*, all insect genomes analyzed have both single genes with high similarity to Orco and multiple genes with similarity to specific ORs. The *M. hrabei* genome did not encode an Orco, but instead contains 5 ORs of high similarity that form a highly supported clade in the gene phylogeny (Figure 2). We also could not find an Orco in the deep RNAseq transcriptome Missbach et al. (2014) generated for their bristletail, *L. y-signata*. This finding leaves open two possibilities. First, bristletails might have lost their Orco gene. Second, the OR family might have originated with a few specific ORs like those of *M. hrabei*, with the Orco lineage evolving between Archaegnatha and Zygentoma. Phylogenetic analysis using various sets of GRs from other insects, arthropods, and animals as outgroup to root the OR family tree does not resolve this question confidently (data not shown). In any case, these five specific ORs in *M. hrabei*, at least one of which is present in *L. y-signata*, must function in the absence of Orco, perhaps alone or as dimers.

Finally, we note that while insects are defined by morphological and developmental synapomorphies (shared derived characters that are unique to a taxon), to the best of our knowledge presence of the OR gene family is the first molecular synapomorphy for the Class Insecta.

## Materials and Methods

A detailed Materials and Methods section can be found in the Supplementary Material.

### Sequencing and assembly of the *Thermobia domestica* genome

Sequencing and assembly of the *T. domestica* genome followed Brand et al. (2018). Briefly, we sequenced the DNA extracted from a single *T. domestica* individual for assembly with the DISCOVAR v1 pipeline (Weisenfeld et al., 2014). Quality assessment of the resulting assembly was based on standard N statistics, k-mer distribution analysis, and the BUSCO v2 pipeline (Simão et al., 2015) as previously described (Brand et al., 2017).

### Odorant receptor annotation

The genomes of the dipluran *Catajapyx aquinolaris*, the collembolans *Holacanthella duospinosa, Orchesella cincta*, and *Folsomia candida*, the firebrat *Thermobia domestica*, the bristletail *Machilis hrabei*, the dragonfly *Ladona fulva*, and the mayfly *Ephemera danica* were used for manual OR gene annotation (detailed in Supplementary Material).

### Odorant receptor gene family analysis

OR protein alignments were produced with CLUSTALX v2 (Larkin et al., 2007) and trimmed using TrimAl (Capella-Gutiérrez et al., 2009). The resulting alignment was used for gene tree inference using RaxML (Stamatakis et al., 2005) under the JTT + G substitution model with 20 independent ML searches and 1000 bootstrap replicates as previously described (Brand and Ramírez, 2017).

## Acknowledgements

We thank Stephen Richards for permission to examine unpublished genome sequences from the i5k pilot project, and Kimberly Walden for assistance with BLAST searches. This work was funded by a National Science Foundation grant, IOS-1456678, to Brian Johnson and Juan Luis Jurat-Fuentes and a Hatch grant to Brian Johnson (CA-D-ENM 2161-H).

